# Cell-cell interactions and evolution using evolutionary game theory

**DOI:** 10.1101/028746

**Authors:** David Basanta

## Abstract

Cancers arise from genetic abberations but also consistently display high levels of intra-tumor heterogeneity and evolve according to Darwinian dynamics. This makes evolutionary game theory an ideal tool in which to mathematically capture these cell-cell interactions and in which to investigate how they impact evolutionary dynamics. In this chapter we present some examples of how evolutionary game theory can elucidate cancer evolution.

## I. Introduction to game theory in cancer

The genetic origin of cancer initiation is well known, as it is the process of clonal evolution and intra-tumor heterogeneity [1], Those facts show cancer as an evolutionary process where the tumor is an evolving population dynamically changing as increasingly more aggressive phenotypes emerge and become dominant [2], [3].

While the genetic aspects of cancer evolution have received much attention, other key features have been comparatively neglected. Those include the role of the physical microenvironment and the role of cell-cell interactions. Other chapters in this booklet deal with agent-based models such as hybrid discrete-continuum cellular automata. These are sophisticated computational tools that allow us to explicitly incorporate properties of the physical microenvironment [4],

The mathematical tool that we will be describing in this chapter is based on Game Theory (GT). First described by Von Neumann and Morgenstern, GT has been used in sociology and economy. A branch of GT, evolutionary game theory (EGT), was created to study population dynamics where interactions between these populations is considered to be key [5], By concentrating on the interactions we are free to momentarily ignore everything else and to see the impact of those on selection pressure.

Much work in EGT has been devoted to the study of evolutionary equilibria and the properties of replicator equations. Equilibria in EGT is based on the concept of Nash equilibria, developed by the recently deceased John Forbes Nash. In a Nash equilibrium none of the players has an incentive to deviate from their current strategy, even if it is suboptimal as any deviation will result in a decrease in fitness for her/him. In EGT an Evolutionary Stable Strategy (ESS) is a type of Nash Equilibrium in which the composition of the population in terms of phenotypic strategies is stable against changes. This is akin to saying that, if we know all the relevant phenotypes, a tumour population in ESS is stable to changes to its current composition. Thus new mutations leading to different phenotypes cannot change the clonal composition of the tumor.

Replicator equations describe the dynamics over time of a game assuming we know the initial composition of phenotypic strategies in the tumour. Together with the ESS this allows us to understand what mutations/phenotypic strategies can emerge in a tumour and the context that can help them be successful.

## II. Using EGT in Cancer

We will illustrate these concepts and how they can be used to shed light on cancer evolution by investigating the emergence of motility and invasiveness in a tumour.

### A. Emergence of invasiveness

Tumours usually start as highly proliferative cells capable of autonomous growth [6], The emergence of invasive phenotypes is thus a worrying but commonly observed event during carcinogenesis. A very simple game would include purely proliferative cells as well as those also capable of motility. This motility comes at a cost (*c*) since cells that are motile at a given time cannot proliferate. Motility can be advantageous though, as it allows a cell to explore new locations that might have more resources (*V*=space/nutrients/oxygen). This can be described with a payoff table like the one in table 1 (top left corner in white). The equilibria can be described by the following (assuming *V*=l):

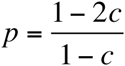

where *p* is the proportion of invasive cells. This equation shows how changing the cost of motility influences the presence of motile cells in the steady state.

**Table 1.**
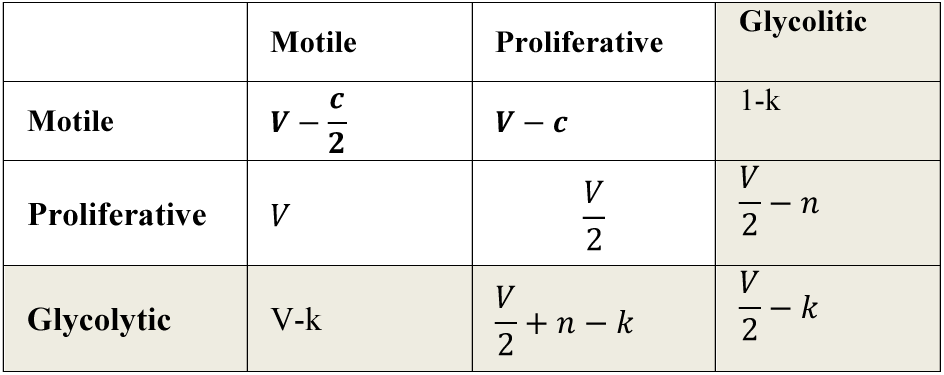
The table shows the change in fitness for cells when interaction with other cells. Thus motile cells interacting with other motile cells do not have to share resources (as one of them will move to a different location). Proliferative cells interacting with other proliferative cells on the other hand have to share existing resources (V).

An extended version of this game (including the shaded bottom-right corner of table 1) allowed us to study how the dynamics of the game change when considering a common type of phenotype found in most types of cancer. Glycolytic cells, that use anaerobic metabolism even in normoxic conditions. This is known as the Warburg effect [7], Interestingly, the new phenotype does not dominate the tumour population but facilitates the emergence of invasiveness by incentivizing motility. Fig. 1 shows this effect on the proliferative population. This suggests that hypoxia could, indirectly, lead to more invasive tumours. The polyclonal equilibria is shown here:

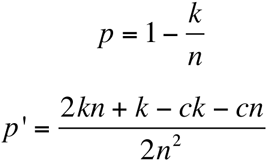

where *p*’ is the proportion of proliferative cells. The ESS shows that an increase in the cost of living in an acid environment and a decrease in the fitness cost of relying on glycolysis leads to more invasive cells, not more glycolytic cells.

**Figure 1.**
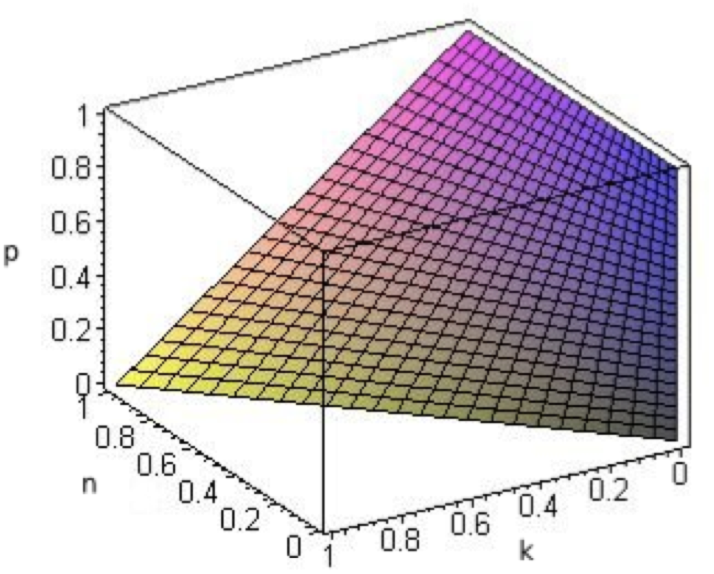
Proportion of invasive cells, *n* is the cost of living in acidity and *k* the cost of a glycolytic metabolism. The proportion of invasive cells increase as we increase the cost of living in an acid environment and decrease the cost of glycolysis, factors that should ease the emergent of glycolytic cells.

### B. Role of space

Mathematical models need to make simplifications. In this model we assumed that space does not need to be explicitly modelled. Removing this simplification allows us to understand how heterogeneously allocated resources and neighbours could impact the game. Recently [8] we used the Othsuki-Nowak transform to investigate how changing the number of other cells a tumour cell will interact with would influence game dynamics. We envisioned that this approach would allow us to understand the different evolutionary dynamics and selection pressures that characterise tumour cells when they reach a boundary or a hard edge. Fig. 2 shows that as cells hit a hard edge, the tumours tend to include more invasive phenotypes and become more polyclonal.

**Figure 2.**
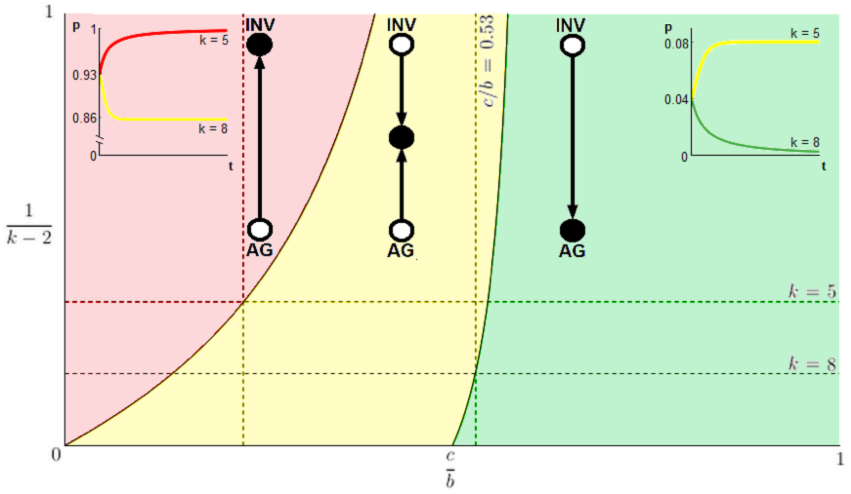
Change in game dynamics as we change number of neighbours. 1/*k* -2 represents the level of viscosity and c/b the relative cost of motility. In the red region the tumour evolves towards motility, in the yellow to coexistence and in green towards exclusively proliferative tumours.

We have also used a Cellular Automaton (CA) *[9]* approach to compare the results of the initial model (excluding glycolytic phenotypes). While it is possible to transform a conventional game into an agent-based model and keep costs and benefits relatively abstract, we built this CA from scratch using the rules shown in fig. 3A. This model recreates the model without relying on the principles of game theory. The results (in fig. 3 B) show the differences between the spatially-explicit model simulations with the EGT-derived analysis. This shows that even when studying tumour invasion, spatially-implicit EGT models can provide useful and accurate qualitative insights.

**Figure 3.**
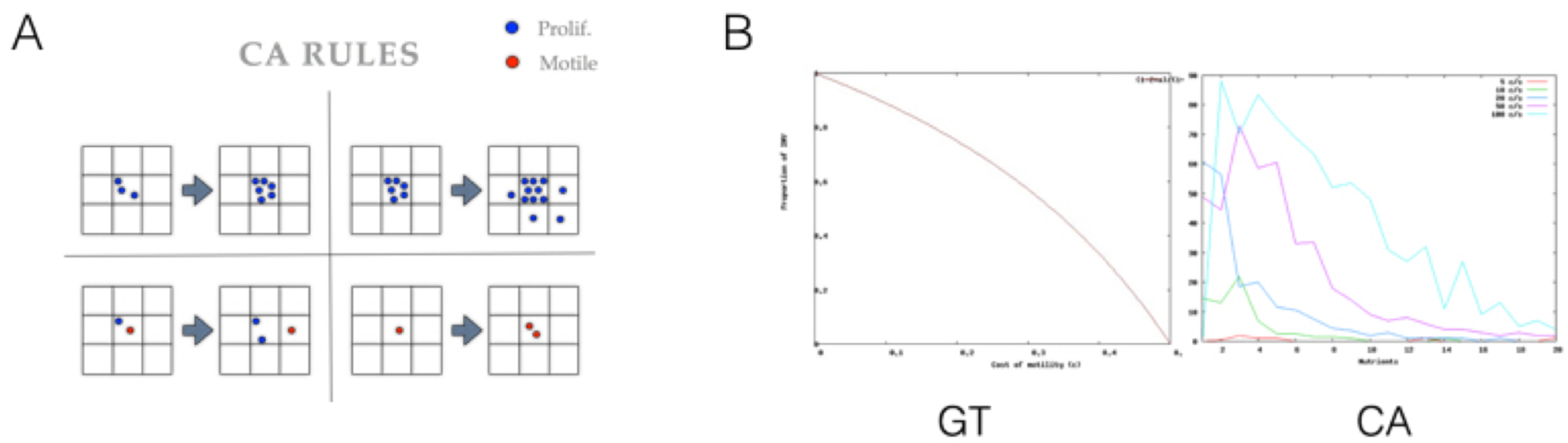
An agent-based model of the invasion game. (A) rules of the agent-based model designed to replicate the proliferation/motility described previously. Motile cells tend to divide when they do not have to share resources. Proliferative cells only move when there is no more space for division left. (B) comparison between EGT analysis (left curve) and agent-based simulations (right plot).

## Acknowledgments

We would like to acknowledge the State of Florida Bankhead Coley program (5BC01), NCI PSOC (5U54CA143970-05) and ICBP (5U01CA151924-04) grants for their generous support.

